# Geometric constraints in the development of primate extrastriate visual cortex

**DOI:** 10.64898/2026.02.04.703881

**Authors:** Hyunju Kim, Michael J. Arcaro, Nabil Imam

## Abstract

Sensory systems are organized into topographic maps that shape information flow and computation across cortical circuits. Although the mechanisms establishing primary sensory maps are well characterized, how higher-order maps emerge across neocortex is unclear. Because the probability and strength of cortical connections fall off steeply with distance along a folded surface, the geometry of the cortex may be a major factor shaping the organization of higher-order maps. To test this in a well-characterized sensory system, we develop a growth model embedded in the folded surface geometry of the macaque visual cortex, using fMRI-defined V1 retinotopy as the sole functional anchor. Cortical organization emerges through algorithmic growth from primary visual cortex, governed by distance-dependent activity correlations and competition among developing projections. Without imposing areal boundaries, map orientations, or predefined topographic layouts, this process generates multiple retinotopic maps with systematic mirror reversals and smooth gradients that reflect key structural features of fMRI-derived extrastriate maps. Parameters estimated on a population template generalize across individual macaques, while individual cortical geometry accounts for fine-scale map variation around a common retinotopic scaffold. These results suggest that conserved growth rules acting on folded cortical surfaces produce stereotyped higher-order retinotopic organization under minimal explicit specification.

## Introduction

A defining property of neocortex is its organization into orderly topographic maps that preserve spatial continuity, so that nearby points in sensory space are represented by nearby locations on the cortical surface [1–3]. This form of organization is present throughout sensory systems, is conserved across species, and emerges early in development [4–10], suggesting that it is a fundamental feature of cortical wiring and functioning. Topographic maps provide an intrinsic coordinate system that shapes how sensory information propagates across cortex, establishing a spatial framework within which later circuit refinement and functional specializations emerge [3, 11, 12]. While the mechanisms that establish primary cortical maps have been studied extensively [5, 13, 14], it remains unclear how topographic organization is established beyond primary cortex, including the arrangement, orientation, and boundaries of successive maps across higher-order cortex.

Theoretical accounts of cortical organization have argued that such structure can arise from general developmental rules acting on an intrinsically topographic substrate, rather than from detailed explicit specification of areal identity or map orientation [11, 13, 15–17]. In this view, maps are treated as the primary organizational units of cortex, and complex areal layouts emerge through self-organizing processes that preserve local continuity while shaping global structure as it unfolds. Early established topographic maps in primary cortex can therefore act as a stable coordinate system guiding subsequent organization under continuity constraints [11, 18, 19]. From this perspective, mirror-reversing representations that distinguish adjacent maps, where the progression of sensory representations reverse directions at each areal border, are not features explicitly encoded in the genome but natural consequences of maintaining continuous topography as activity-dependent interactions spread across the cortical sheet [11, 13, 16]. A central prediction of this account is that the arrangement and orientation of maps should follow from conserved growth rules operating on the cortical surface without explicit instructions specifying boundaries or reversals.

Self-organizing growth models provide a concrete way to formalize and test these ideas. Prior work has shown that a small set of developmental processes based on distance-dependent interactions and wiring constraints can give rise to sequential mirror-reversing maps [16, 17]. In these models, higher-order map structure emerges through activity-driven growth originating in anchor regions such as the LGN or primary visual cortex, without explicitly specified areal labels or reversal rules. However, these demonstrations were carried out on simplified substrates, typically planar grids with idealized distance relationships, leaving open the question of whether self-organizing principles operating on real cortical surfaces with complex geometries can recapitulate higher-order map organization observed in the brain.

This question is particularly important given that cortical connectivity is strongly distance-dependent, with connection probability and strength falling off steeply with physical distance between cortical locations [20, 21]. Interareal projections are additionally shaped by laminar organization and cortical folding, linking surface geometry to long-range connectivity patterns [22–24]. Moreover, developmental growth proceeds on a folded, anisotropic cortical sheet, where much of the increase in connection length reflects stretch-driven expansion of an existing axonal scaffold rather than late formation of new long-range projections [16, 25]. As a result, the geometry of the cortical surface directly constrains which interactions are most likely to occur during development. This motivates examining whether a developmental process embedded in empirically measured cortical geometry can reproduce key features of retinotopic organization under distance-dependent interactions and wiring constraints alone.

The primate visual system, in which multiple retinotopic maps are distributed across occipital and temporal cortex [26–30], offers an ideal test case for these principles. Beyond primary visual cortex, extrastriate areas traditionally labeled V2, V3, and V4 each contain orderly representations of polar angle and eccentricity arranged in a stereotyped spatial layout [11, 28, 31]. This pattern is remarkably reliable across individuals, despite substantial variability in cortical folding and surface geometry [3, 11, 29, 30]. Importantly, this organization is present at birth [10, 32, 33], indicating that it must arise from early developmental processes prior to extensive visual experience.

Here, we derive a network model of cortical development embedded directly within the geometry of the macaque visual cortex, reconstructed from structural and functional MRI. Cortical organization unfolds as a directed growth process from primary visual cortex, shaped by distance-dependent activity correlations and wiring constraints, consistent with mechanisms of circuit formation observed in the cortex [34, 35]. Without imposing areal boundaries, map orientations, or reversal rules, the model gives rise to multiple orderly retinotopic maps characterized by mirror-reversing phase organization and smooth eccentricity gradients. We apply the same growth rules across population-averaged and individual macaque cortical surfaces to assess the robustness of emergent map structure and to examine how individual differences in cortical geometry across monkeys shape variation around a shared organizational scaffold. These analyses show that stereotyped retinotopic organization of higher-order visual areas can emerge through activity-dependent self-organization under minimal explicit specification.

## Results

### Growth Model Applied to Neocortex

To model cortical map formation under realistic anatomical constraints, developmental dynamics were embedded in the folded surface geometry of the macaque visual cortex. The model was formulated as a directed growth process that unfolds in a network, in which nodes represent localized regions sampled from the cortical sheet, each associated with retinotopic tuning measured using functional MRI [8]. Directed edges represent projections originating from primary visual cortex and terminating within a contiguous extrastriate region of visual cortex. Primary analyses were performed on the NIMH Macaque Template (NMT), a population-averaged macaque cortical reference [36], with model simulations conducted separately for the left hemisphere (LH) and right hemisphere (RH). Combined with group average retinotopy, the NMT provides a common anatomical and functional reference frame that captures the typical spatial arrangement of visual maps across individuals. This offers a principled baseline for parameterization and visualization before assessing how individual cortical geometry gives rise to retinotopic organization in individual animals.

To represent growth across the cortical sheet while preserving geodesic distances defined by the folded surface, cortical locations were embedded into a two-dimensional coordinate system derived from the measured surface distance matrix (Fig. 1A–C, see Methods). This transformation unfolds the three-dimensional cortical sheet into a flat representation in which pairwise distances are maintained, so that nearby cortical locations remain nearby despite cortical folding. Distances measured between locations in the embedded space were therefore highly correlated with geodesic distances computed on the original cortical surface, with a Spearman rank correlation coefficient of *ρ* = 0.99 for both hemispheres. The resulting embedding allows developmental dynamics to operate on a two-dimensional sheet that facilitates analysis and visualization while preserving spatial relationships among cortical locations.

**Figure 1:**
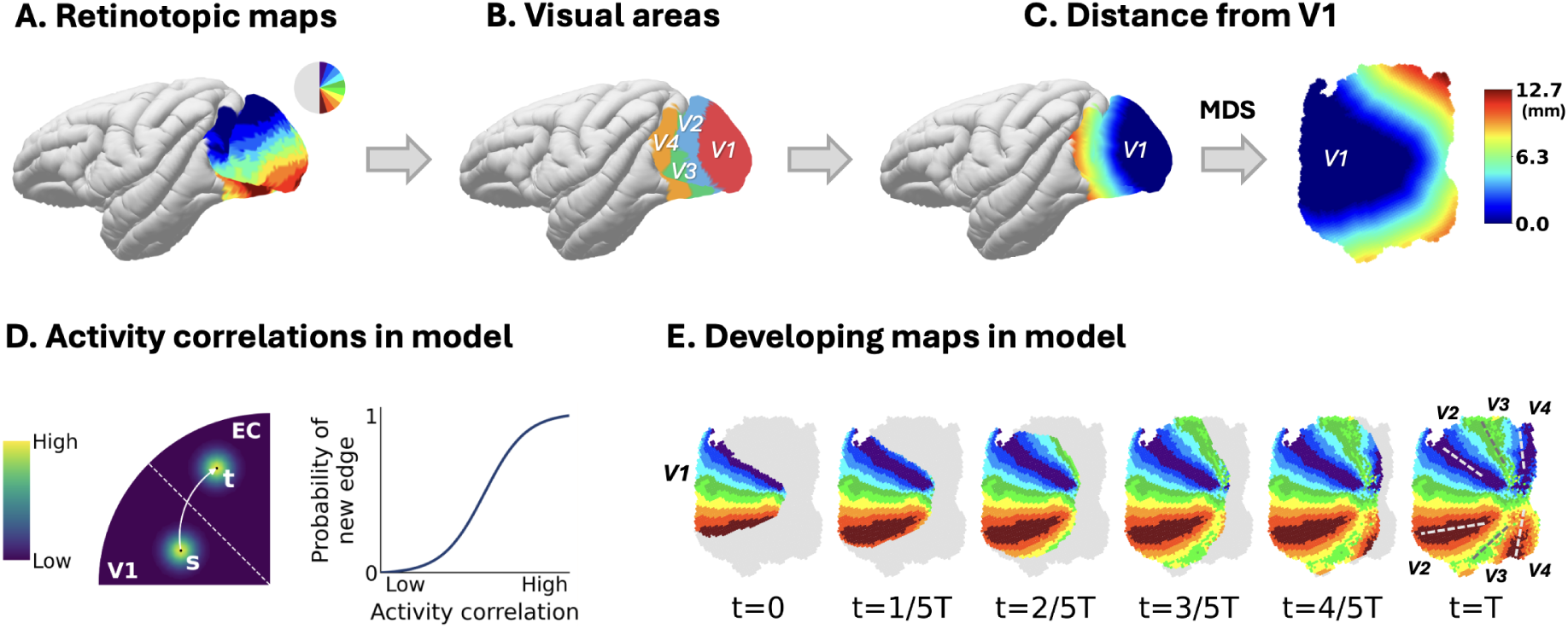
Geometry-constrained growth model of extrastriate cortical maps. **A.** Empirical retinotopic maps in the macaque visual cortex. Colors indicate polar angle representation in contralateral visual space, as shown in the color wheel inset, with red corresponding to the upper vertical meridian, blue to the lower vertical meridian, and green to the horizontal meridian. **B.** Visual areas (V1–V4) defined from the retinotopic map. **C.** Cortical geometry used by the model. Geodesic distance from V1, shown on the cortical surface (left) and after 2D embedding (right) via multidimensional scaling (MDS), which preserves pairwise distances measured along the folded cortical surface. The color bar indicates geodesic distance from V1 in millimeters. **D.** Activity-dependent correlations and edge formation in the model. Left: Activity of a V1 node s generates correlated activity in its spatial neighborhood, as well as in the neighborhood of a distant node t to which s projects. EC denotes extrastriate cortex. Right: probability of forming a new edge as a function of activity correlation between two nodes. **E.** Developing maps in the model. Polar-angle maps on the cortical surface at six time points, where T is the total number of growth steps. Dashed lines at t = T mark the inferred boundaries between higher visual areas V2, V3, and V4, identified from the final polar-angle map. The areas emerge naturally from activity correlations and competitive growth.

Cortical map formation was modeled as a process of algorithmic growth, in which directed edges (connections) were progressively added from primary visual cortex to nodes within the extrastriate cortical territory. At each step, candidate edges were evaluated using a growth rule that is based on distance-dependent activity correlations and competition among developing projections (Fig. 1D, see Methods). Activity correlations drive the emergence of similar receptive fields among coactive nodes, consistent with Hebbian principles, whereas the competition term reduces the likelihood that the same primary visual cortex nodes repeatedly form new connections. As edges are added, both activity correlations and the competition term are updated, dynamically modifying pairwise node affinities that govern subsequent edge selection. This interaction between distance-dependent correlations and competition for outgoing connections defines the model mechanism linking cortical surface geometry to emergent retinotopic organization beyond V1. The model consists of only two free parameters defining the spatial extent of activity correlations along orthogonal directions of the cortical surface.

### Emergent Retinotopic Topography in the Template Brain

To assess whether growth constrained by cortical surface geometry is sufficient to generate higher-order retinotopic organization, the model was initiated from primary visual cortex and allowed to unfold across the cortical surface. As the process unfolded, extrastriate cortex developed an emergent large-scale topographic organization, with retinotopic representations varying smoothly across the cortical sheet (Fig. 1E). This organization arose incrementally as growth propagated away from V1, demonstrating that surface-constrained growth dynamics can give rise to structured retinotopic layouts beyond primary visual cortex.

Across extrastriate cortex, the retinotopic maps generated by the model recapitulated key features of macaque retinotopy spanning areas V2, V3, and V4 as measured with fMRI. In model predictions, representations of the lower visual field occupied dorsal extrastriate cortex, while upper visual field representations were located ventrally, reflecting the characteristic inversion of visual space observed in macaque visual cortex (Fig. 2A,B). Eccentricity organization showed a smooth progression away from the foveal representation, oriented roughly orthogonal to polar angle organization and forming a continuous large-scale gradient that extends across extrastriate cortex, in contrast to the mirror reversals observed for polar angle. Quantitatively, retinotopic organization in extrastriate cortex showed strong agreement between model predictions and fMRI measurements (Fig. 2A,B), with polar angle maps exhibiting strong rank-order correlation (Spearman’s *ρ* = 0.87 LH; 0.84 RH) and eccentricity maps showing moderate-to-strong correlation (LH: *ρ* = 0.65; RH: *ρ* = 0.64). Together, these results show that retinotopic correspondence emerges without explicit specification of visual field layout or spatial gradients, arising instead from a small set of developmental rules acting on the inherited topographic structure of primary visual cortex.

**Figure 2:**
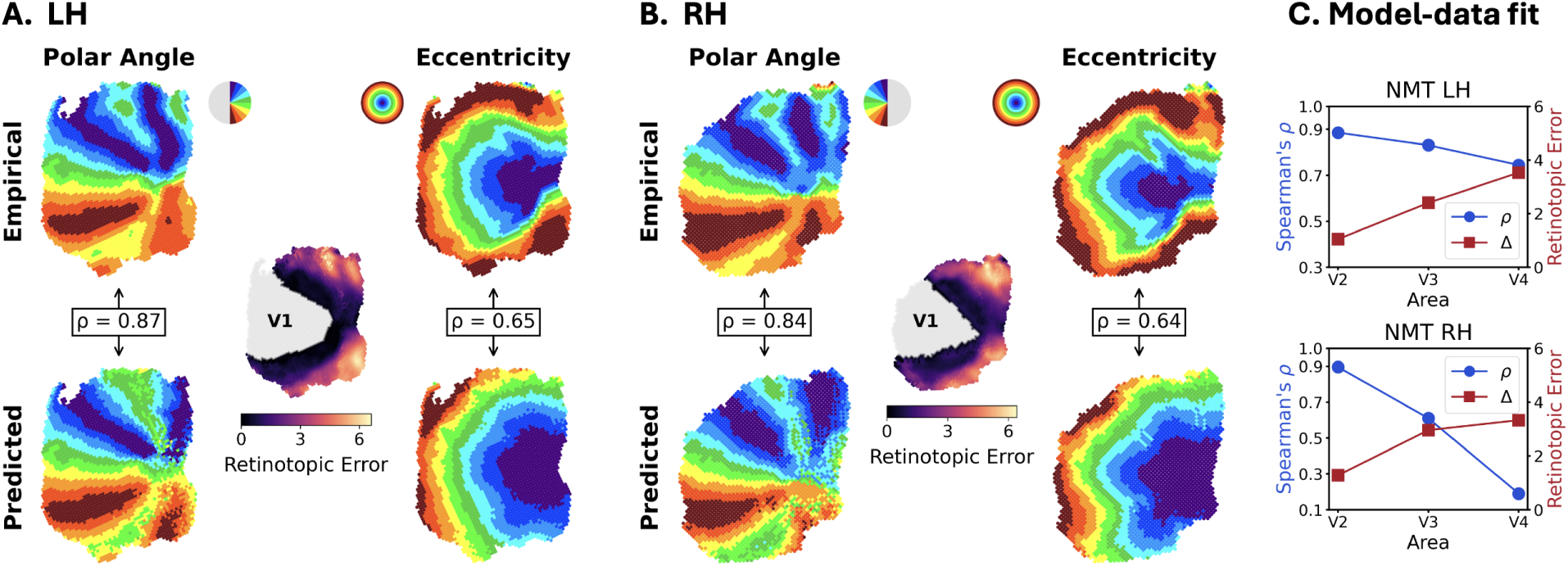
Empirical vs. predicted retinotopy in NMT. **A.** Left hemisphere (LH) polar angle and eccentricity maps (empirical, top; predicted, bottom). Spearman’s ρ computed across extrastriate nodes (V2–V4; excluding V1) is ρ = 0.87 for polar angle and ρ = 0.65 for eccentricity. The middle inset shows node-wise retinotopic error Δ in visual degrees (Eq. 5), computed as the Euclidean distance between predicted and empirical retinotopic coordinates. **B.** Same as **A** for the right hemisphere (RH; ρ = 0.84 for polar angle; ρ = 0.64 for eccentricity). **C.** Area-wise summary across extrastriate cortex showing mean retinotopic error and retinotopic correspondence (mean Spearman correlation of polar angle and eccentricity), averaged across nodes within each area for V2–V4 (top: LH, bottom: RH).

Overall retinotopic error remained low across cortical locations, with error defined as the distance between predicted and empirical coordinates in retinotopic space, expressed in degrees of visual angle (^◦^). Area-wise analysis revealed a gradual decrease in retinotopic correspondence from V2 to V4, accompanied by increasing error (Fig. 2C). In NMT LH, mean Spearman correlation was 0.89 in V2, 0.83 in V3, and 0.74 in V4, with corresponding retinotopic errors of 1.05^◦^, 2.41^◦^, and 3.53^◦^, respectively. In NMT RH, mean Spearman correlation was 0.90 in V2, 0.61 in V3, and 0.19 in V4, with retinotopic errors of 1.28^◦^, 2.96^◦^, and 3.33^◦^, respectively. The largest disagreements were concentrated in peripheral portions of ventral V4, where the model tended to produce representations shifted toward more parafoveal locations. This pattern suggests a distance-dependent influence of the V1 anchor: with increasing distance from V1, the organization of extrastriate cortex is less directly constrained by the V1 reference frame, allowing small deviations relative to that anchor to accumulate and appear as larger differences between predicted and empirical retinotopic coordinates. Consistent with this interpretation, empirical work has shown that covariance between the sizes of V1 and extrastriate visual maps decreases with increasing cortical distance from V1 [37]. Together, these results show that growth constrained by realistic cortical-surface geometry is sufficient to reproduce the large-scale spatial layout of retinotopic organization observed in macaque extrastriate visual cortex.

To assess whether the observed correspondence reflects a specific alignment of retinotopic organization rather than a generic consequence of smooth topographic structure, we examined the effects of systematically rotating the predicted tuning phase maps prior to comparison with empirical fMRI retinotopy (Fig. 3). Correspondence in the spatial arrangement of polar angle organization was strongest without rotation. Quantitatively, the reference alignment (0^◦^) produced the highest correspondence and lowest retinotopic error across hemispheres (LH: *ρ* = 0.87, Δ = 2.10^◦^; RH: *ρ* = 0.85, Δ = 2.22^◦^). Correspondence declined progressively with increasing rotation, with reduced alignment at 45^◦^ (LH: *ρ* = 0.41, Δ = 5.57^◦^; RH: *ρ* = 0.40, Δ = 3.20^◦^), reaching the poorest agreement at 90^◦^ (LH: *ρ* = −0.43, Δ = 7.97^◦^; RH: *ρ* = −0.81, Δ = 4.53^◦^), and partially recovering at 135^◦^ (LH: *ρ* = 0.49, Δ = 7.29^◦^; RH: *ρ* = 0.20, Δ = 3.57^◦^). This pattern indicates that the growth model captures specific large-scale alignment of retinotopic organization observed in macaque visual cortex, rather than merely producing generic smooth topographic structure.

**Figure 3:**
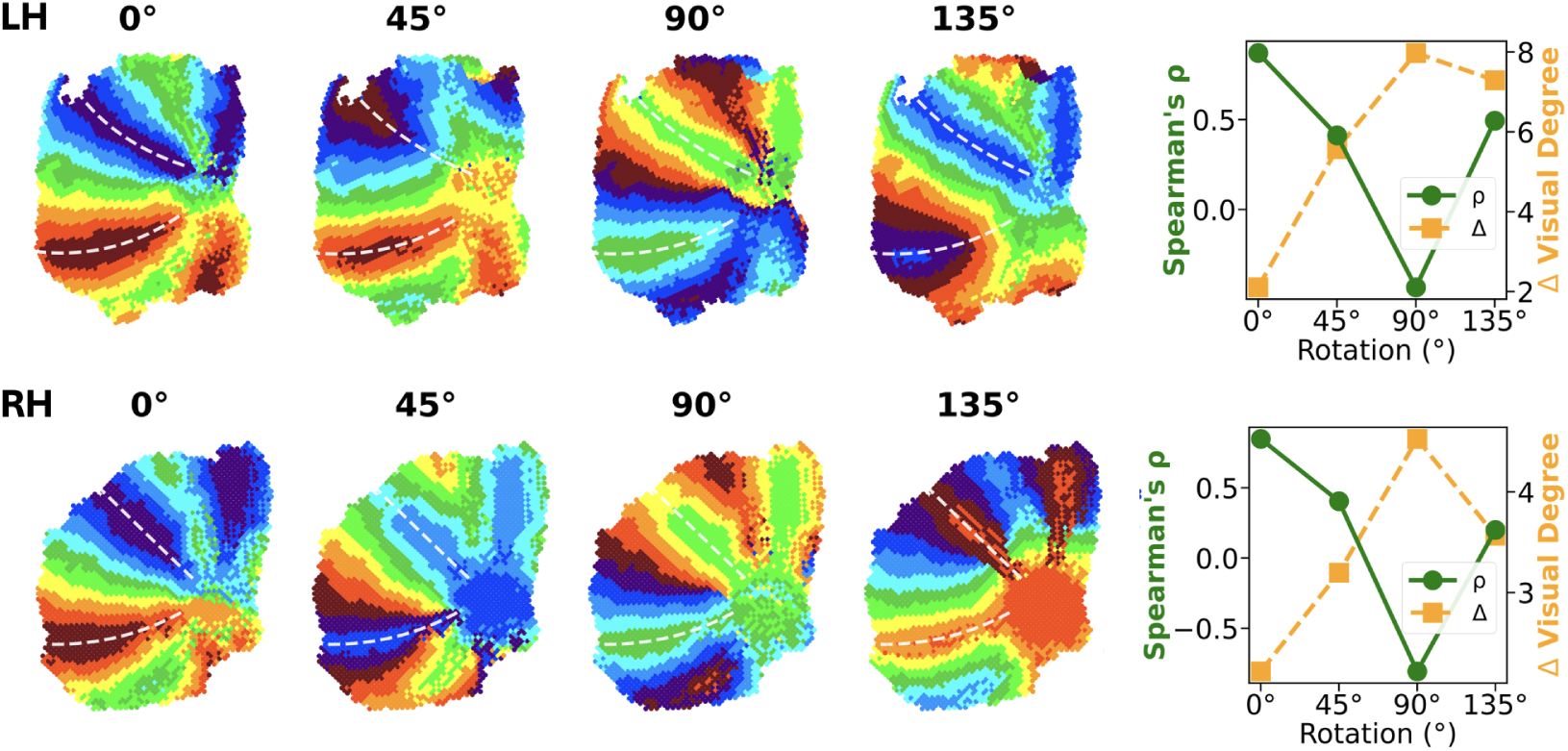
Rotation control for global polar angle phase alignment (top: NMT LH, bottom: NMT RH). Retinotopic phase maps are shown for model predictions after systematically rotating the V1 polar angle phase map (0^◦^, 45^◦^, 90^◦^, 135^◦^) while keeping the growth model and its V1-to-extrastriate connections fixed. For each rotation, predicted extrastriate phase maps were recomputed by propagating the rotated V1 phase values through the model projections, yielding a family of alternative global phase alignments. Quantitative correspondence with empirically measured retinotopy is maximal for the unrotated configuration. Summary metrics show mean Spearman rank correlation (ρ; polar angle) and mean retinotopic error (Δ) as a function of rotation angle, averaged within each hemisphere. A 90^◦^ rotation, corresponding to half of the tuning space, yields the poorest agreement.

The emergent retinotopic organization was robust to variations in model parameterization and implementation. The model consists of a total of two free parameters, *σ*_R_ and *σ*_T_, that define the spatial extent of activity correlations (see Methods). A grid search over these two parameters revealed a broad region of parameter space yielding stable retinotopic organization (Supplementary Figs. S1–S3; see Supplementary Materials, Secs. A–C). Retinotopic structure was also preserved across alternative node incorporation orders during growth, including periphery→ fovea, fovea→ periphery, and random sequences (Supplementary Fig. S4; see Supplementary Materials, Sec. D), indicating that map structure is robust to eccentricity-dependent developmental ordering, which might have been expected to affect map formation given the protracted development of the fovea [38] and the earlier maturation of the magnocellular pathway [39].

In the model described so far, retinotopic maps form through projections exclusively from V1 to extrastriate cortex, consistent with theories in which early-established primary areas organize cortical maps [11, 16, 18, 19]. This setup does not assume explicitly defined developmental stages across a processing hierarchy (V1→V2→V3→V4). To assess if hierarchical development improves the match between model and data, we evaluated a variant of the model in which projections unfolded sequentially across successive source populations rather than arising from V1 alone (see Supplementary Materials, Sec. E). This formulation required additional specification of developmental timing, which enforced V1→V2 connections to establish first before allowing V2 to project edges, and so on for V2→V3 and V3→V4. Under this formulation, the model produced retinotopic organization closely matching the results of our primary model (Fig. S5A,B), indicating that explicitly staged hierarchical growth is not necessary for the emergence of retinotopic map structure.

### Mirror-reversing Retinotopic Maps

In primate visual cortex, boundaries between adjacent extrastriate visual areas are defined by reversals in the progression of polar angle representations along the cortical surface. These reversals mark transitions between distinct retinotopic maps within a continuous topographic organization and provide a functional criterion for identifying map structure independent of anatomical labels.

Consistent with this organizational principle, the emergent topography produced by the growth model exhibited systematic mirror reversals in polar angle representation across extrastriate cortex (Fig. 4). As growth propagated away from primary visual cortex, polar angle representations progressed smoothly over extended cortical regions and then reversed direction at consistent anatomical locations within extrastriate cortex, aligned along the principal axis of propagation. This pattern produced an alternating sequence of polar angle progressions characteristic of multiple retinotopic maps. Notably, these polar angle reversals were not imposed by predefined areal boundaries, map orientations, or reversal rules. Instead, they emerged because the correlation driven growth rule promotes coherent structure both within local extrastriate neighborhoods and in the mapping from V1 inputs onto extrastriate targets. Thus, mirror reversals reflect an emergent property of maintaining smooth topography under surface constrained growth, rather than an explicitly specified feature of the model.

**Figure 4:**
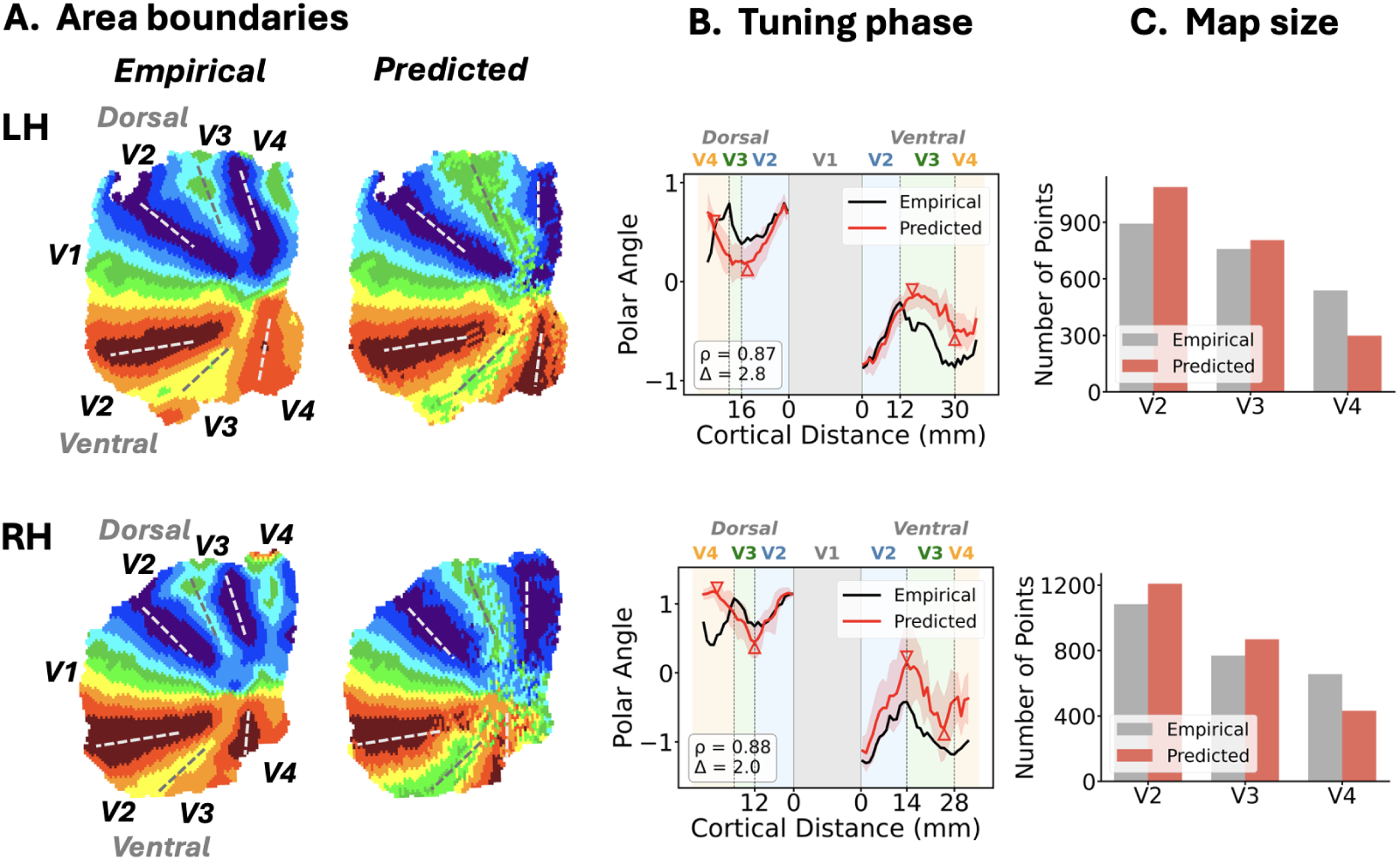
Mirror-reversing retinotopic phase organization and derived visual area boundaries (top: NMT LH, bottom: NMT RH). **A.** Polar angle phase maps for empirically measured macaque fMRI retinotopy and model-predicted organization are shown for each hemisphere, together with inferred boundary locations separating successive mirror-reversing retinotopic maps. Dashed white lines indicate reversals at the vertical meridian, whereas dashed grey lines indicate reversals at the horizontal meridian. **B.** Polar angle tuning along geodesic distance profiles extending away from the V1 border in two opposite directions. The direction extending toward the left side of the profile corresponds to lower visual field representations (dorsal stream), whereas the direction extending toward the right side corresponds to upper visual field representations (ventral stream). Black and red curves denote empirical and predicted tuning, respectively, with shaded regions indicating across-subject variability. Colored bands at the top mark the extent of each visual area. Spearman’s ρ and mean absolute difference Δ between empirical and predicted curves are reported. **C.** Number of cortical points assigned to extrastriate areas V2, V3, and V4 in the empirical (gray) and predicted (red) maps, providing a direct comparison of relative surface area.

To quantify the organization of polar angle representations and compare model predictions with empirical macaque retinotopy, we summarized polar angle structure using a phase–distance anal-ysis applied to both predicted and fMRI-derived maps. For each hemisphere, retinotopic tuning coordinates were expressed in a V1-anchored polar coordinate system, and nodes were grouped into eccentricity bands based on V1-referenced eccentricity. Within each band, nodes were assigned a signed geodesic distance from the V1 border, separately for dorsal and ventral directions, and polar angle phase values were averaged within distance bins to obtain one-dimensional phase–distance profiles (Fig. 4A). The resulting profiles exhibited clear periodic structure, with alternating increases and decreases in phase marking reversals in polar angle progression along the cortical surface. These reversal points correspond to transitions between adjacent retinotopic maps and provide operational markers of boundaries between extrastriate areas (Fig. 4B,C).

Across hemispheres, phase–distance profiles derived from the growth model were consistent with those obtained from macaque fMRI data. In both predicted and empirical maps, phase reversals occurred at similar cortical distances from the V1 border along both dorsal and ventral trajectories (Fig. 4B,C), with a small deviation at the dorsal V3–V4 boundary, which appeared slightly anterior in the model. Consistent with this qualitative agreement, predicted and empirical profiles showed strong correspondence, with high Spearman correlations (LH: *ρ* = 0.87, RH: *ρ* = 0.88) and small mean phase differences (LH: 2.8^◦^, RH: 2.0^◦^). Because these reversal locations define functional boundaries between adjacent retinotopic maps, they were used to estimate the relative spatial extent of areas V2, V3, and V4. Model-derived area extents captured both the relative sizes of individual areas and the systematic decrease in surface area across successive extrastriate maps with increasing distance from V1, as observed in the fMRI data.

### Geometry Constrains Individual Retinotopy

Having established that the growth model reproduces the stereotypical large-scale retinotopic organization observed in macaque visual cortex, the next question was whether differences in cortical geometry further constrain retinotopic maps at the level of individual monkeys. To address this, the growth model was applied independently to each macaque dataset using its native cortical surface geometry, while all growth rules and parameters derived from the template analysis were held fixed across animals. Model predictions were then compared directly with empirically measured retinotopic maps from the same monkeys, allowing evaluation of whether individual cortical folding and surface layout carry information that shapes individual retinotopic organization beyond the population-averaged template.

Across monkeys, the growth model captured features of the individual retinotopic organization observed in the empirical fMRI data (Fig. 5A, Fig. S6; Supplementary Materials, Sec. F). In the measured maps, extrastriate visual cortex showed orderly progression of polar angle along the cortical surface, with consistent reversals marking transitions between adjacent retinotopic maps. Predicted eccentricity maps exhibited smooth gradients extending away from the foveal representation that closely aligned with those measured using fMRI. Specifically, Spearman correlations between model-predicted and fMRI-derived maps, computed separately for the left and right hemispheres and then averaged within each monkey, were consistently high across monkeys (0.74 ± 0.03 for polar angle and 0.60 ± 0.10 for eccentricity). Together, these results demonstrate that both angular and radial components of retinotopic organization emerge robustly on individual cortical surfaces and reflect individual deviations beyond a shared, species-typical retinotopic scaffold.

**Figure 5:**
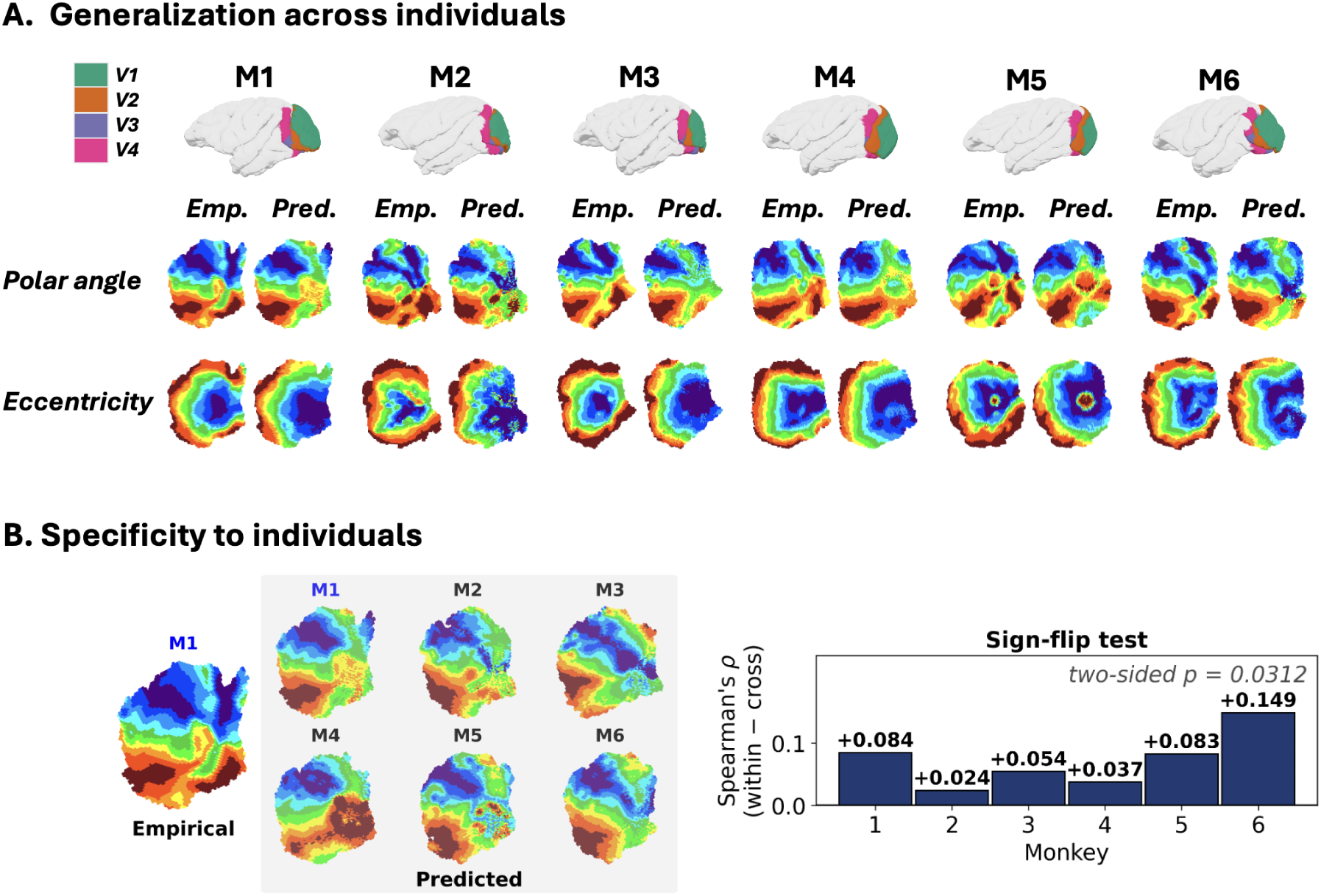
Conserved growth rules and individual cortical geometry jointly determine retinotopic organization. **A.** The same growth rules and model parameters generalizes across all six monkeys (M1–M6). Shown are empirical (Emp.) and predicted (Pred.) polar angle and eccentricity maps for the left hemisphere of each monkey, alongside their cortical surfaces (top row). **B.** Variation of fine-scale retinotopy is specific to each individual. Left: Extrastriate polar angle maps for monkey M1 predicted using V1 tunings from each monkey (M1–M6). M1’s empirical map is also shown for comparison. Coarse retinotopic structure is preserved under tuning transfer, but fine-scale alignment degrades. Right: Per-monkey difference in Spearman’s ρ between within-monkey and cross-monkey predictions. All six monkeys show a positive value (two-sided sign-flip test, p = 0.0312), indicating that an individual’s specific cortical folding pattern shapes its V1 to extrastriate mapping.

To evaluate whether accurate prediction of individual retinotopic maps depends on the alignment between cortical geometry and V1 tuning statistics, we performed a complementary analysis in which the growth model was run on each individual macaque cortical surface while substituting V1 tuning statistics from other animals (Fig. 5B). For a given monkey, cortical surface geometry and growth parameters were held constant, and only the retinotopic tuning of V1 inputs was exchanged across animals. When tuning statistics from other monkeys were used, the model continued to produce coherent retinotopic organization, indicating that global map structure is strongly constrained by cortical geometry and growth dynamics. However, systematic deviations emerged in fine-scale phase alignment relative to the empirical maps. Correspondence with empirical retinotopy tended to be higher for within-monkey predictions, with all monkeys exhibiting a positive within-monkey minus cross-monkey difference, a positive mean difference across monkeys, and a reliable bias toward higher within-monkey correspondence. The latter was confirmed by an exact two-sided sign-flip test that enumerates all 2^6^ possible sign assignments under the null hypothesis of zero mean advantage (*p* = 0.0312). Together, these results indicate that the structure of extrastriate retinotopic organization is strongly shaped by geometric features shared across macaque cortices, while individual differences in cortical geometry systematically modulate this structure to produce monkey-specific connectivity patterns.

## Discussion

We show that higher-order retinotopic organization can emerge from growth dynamics constrained by realistic cortical surface geometry. When embedded within folded macaque cortical anatomy and initiated from primary visual cortex, a distance-dependent growth process generated smooth retinotopic gradients, systematic mirror reversals in polar angle, and spatial layouts that closely match retinotopic organization measured using macaque fMRI. These results demonstrate that large-scale extrastriate map organization, including the relative placement of multiple retinotopic maps, can arise without explicit specification of visual area boundaries, map orientations, or reversal locations. Instead, this pattern follows from conserved growth rules acting on the folded cortical sheet together with inherited topographic structure from primary visual cortex.

### Primary Cortical Anchors and Emergent Map Organization

These results support theoret-ical accounts in which early established topographic maps in primary cortex act as developmental anchors for subsequent cortical organization. In primates, primary visual cortex provides a stable retinotopic coordinate system that constrains the emergence of higher-order maps through continuity-preserving growth dynamics [3, 11]. Within this framework, mirror-reversing representations are not imposed markers of areal identity but arise naturally from extending continuous topography across the cortical surface under distance-constrained interactions [16]. The present model formalizes this idea within measured cortical surface geometry, demonstrating that mirror reversals emerge from correlation-driven growth and competition among developing projections. Together, these factors generate coherent structure within extrastriate neighborhoods, defined by the mapping from primary visual cortex onto extrastriate targets, without requiring explicit boundaries or reversal rules.

Prior modeling work has shown that continuity and competition are, in principle, sufficient to generate higher-order retinotopic structure. Broadly, these models fall into two categories: algorithmic growth models, in which mirror-reversing maps emerge from developmental wiring rules [16, 17], and optimization-based models, in which higher-order spatial structure arises once model parameters are optimized for functional objectives [40, 41]. Both approaches, however, have either assumed idealized geometry or fixed areal layouts, leaving unresolved how real cortical anatomy shapes the emergence and placement of retinotopic maps across cortex. The present results address this gap by showing that cortical geometry plays an active role in shaping continuity-based organization, constraining the emergence and placement of retinotopic maps across cortex. When growth dynamics are embedded in measured macaque cortical surfaces, the resulting retinotopic maps reproduce the spatial layout observed with fMRI, with individual differences in cortical folding predicting individual differences in map arrangement around a shared organizational scaffold.

The anchoring framework also highlights important limits. When multiple anchors converge within the same cortical territory, continuity constraints may conflict, leading to local discontinuities in topographic structure [11,42]. In addition, the influence of primary sensory anchors may weaken with increasing distance from early sensory cortex, allowing greater flexibility in higher-order regions [43]. The present model supports this view—correspondence between predicted and empirically measured retinotopic organization systematically decreased with increasing distance from V1, as reflected by rising prediction error from V2 to V3 to V4. This pattern parallels prior empirical findings showing a decrease in spatial correlation between extrastriate visual areas and V1 at increasing cortical distances [37]. Together, these results suggest that early extrastriate organization remains strongly anchored to primary cortex, whereas reduced anchoring at greater distances may permit increased variability in areal size, layout, and functional specialization. Extending the present model to incorporate multiple anchors or pathway-specific growth rules may therefore be necessary to explain organization beyond early extrastriate areas tested in the current study, particularly where retinotopic maps diverge across parallel processing streams.

### Generalization across Individuals and Species

A central result of this study is that the same growth rules and parameters reproduce retinotopic organization across individual macaques when applied to each animal’s native cortical geometry. Despite inter-individual variability in cortical folding and surface layout, the model captures the spatial arrangement of retinotopic maps observed in each monkey, indicating that individual differences in cortical geometry constrain how a shared species-typical scaffold is expressed [8, 26].

Evaluating whether these principles extend beyond macaques is important for two related reasons. First, the substantial differences in scale and systems-level organization between human and macaque visual cortex provide a strong test of whether shared geometry-constrained growth rules can generate distinct map organizations across species. Second, the successful application of the same model to multiple primate species carries implications for developmental timing. Large-scale resting-state fMRI studies show that human visual cortex is already organized at birth into distinct pathways with posterior-to-anterior hierarchical structure and adult-like topographic organization, and that this organization strengthens across late gestation [44]. In macaques, retinotopic proto-organization is likewise present at or near birth [8]. If growth dynamics can account for map organization in both humans and macaques, it would explain why retinotopic organization emerges early in both species prior to extensive postnatal experience.

Cross-species comparisons also clarify which aspects of retinotopic organization generalize as the cortex expands. Continuity-based models suggest that increasing cortical surface area can yield additional mirror-reversing maps as cortex is filled in under smoothness constraints [16]. This is motivated by empirical work proposing a roughly 10-fold increase in the number of cortical areas from early mammals to primates [15, 45]. However, in primates, cortical surface expansion does not necessarily translate into a greater number of early retinotopic maps. Recent work comparing macaques and humans argues that primate visual cortex evolution is dominated by expansion of a conserved architectural scaffold rather than proliferation of new early maps [37]. Applying geometry-constrained growth models to human cortical surfaces therefore provides a direct test of whether conserved developmental rules predict conserved map number while allowing species differences to emerge in map spacing and pathway organization, consistent with network-level perspectives emphasizing distributed and interacting pathways [46].

### Developmental Timing and Anatomical Constraints

The growth model used here was embedded in cortical surfaces derived from adult macaque brains, even though retinotopic maps emerge very early in development. In macaques, large-scale retinotopic organization is already present at or near birth, with orderly polar angle and eccentricity gradients observed across visual cortex prior to extensive visual experience [8]. Although the cortex expands substantially after birth, available evidence suggests that this expansion preserves the relative spatial organization of visual cortex. Despite large increases in cortical surface area across development, the proportional arrangement of visual areas relative to primary visual cortex remains stable [37]. These observations support the use of adult cortical surfaces as a reasonable approximation of the anatomical constraints relevant for early map formation, while motivating future work embedding the model in fetal or neonatal cortical reconstructions.

### Limitations and Future Work

Several directions remain open for extending the presented framework. The growth model was embedded in cortical geometries derived from adult macaque brains, even though retinotopic maps emerge very early in development, with orderly polar angle and eccentricity gradients present at or near birth prior to extensive visual experience [8]. Although the cortex expands substantially after birth, available evidence suggests that this expansion preserves the relative spatial organization of visual cortex [37], supporting the use of adult cortical geometry as a reasonable approximation of the anatomical constraints relevant for early map formation, while motivating future work embedding the model in fetal or neonatal cortical reconstructions. A related direction concerns how interaction structure is defined in the model. Activity correlations were parameterized using distances measured along the cortical surface, which capture key spatial constraints imposed by the folded cortical sheet; however, true axonal path lengths traverse white matter and may diverge from surface geodesic distance. Moreover, cortical folding itself likely reflects developmental constraints imposed by connectivity rather than serving as a purely independent substrate [25]. While surface distance is therefore a reasonable proxy for relative interaction strength across much of visual cortex, especially for short- and intermediate-range connections, exploring alternative distance formulations, including unfolded geometries or tract-informed measures, would help clarify which aspects of cortical geometry are most critical for map spacing and layout.

The model also treats cortical nodes as uniform units, despite known variation in neuronal density, receptive field size, and representational scale across visual areas. While such factors may contribute to quantitative differences in map spacing and transition sharpness along the visual hierarchy, the present results show that the large-scale organization of retinotopic maps can arise without assuming such area-specific properties. Future work could also consider additional cortical anchors beyond V1. Area MT receives direct LGN input and has been proposed as a secondary anchor for dorsal visual cortex [11]. Incorporating such an anchor could introduce competitive constraints on developing projections that influence areal boundaries, potentially accounting for the slightly anterior placement of the dorsal V3/V4 border in the model relative to empirical data.

### Implications for Cortical Area Formation

More broadly, these results support theories of cortical map formation in which areas emerge as stable patterns within a continuous topographic field rather than as discrete units requiring explicit specification of areal boundaries [3,11,16]. In this framework, mirror-reversing structure reflects transitions in map orientation that arise naturally from continuity-preserving dynamics on a folded surface, and map boundaries can be defined operationally from intrinsic organization rather than from predefined labels. Building on this framework, the present model establishes a mechanistic link between realistic cortical geometry and the emergence and spatial arrangement of multiple retinotopic maps. Together with recent accounts of functional organization at multiple stages of the visual hierarchy [40, 41], these results suggest that wiring, geometry, and growth dynamics jointly shape how cortical maps emerge, stabilize, and diversify across individuals and species.

## Methods

### Data Acquisition

Retinotopic mapping data were acquired from six macaque monkeys using a 3T TimTrio scanner with a 4-channel surface coil at the Athinoula A. Martinos Center for Biomedical Imaging, Massachusetts General Hospital. Functional images were collected using an EPI sequence at 1 mm isotropic resolution. High-resolution anatomical images were acquired with a 3T Siemens Skyra scanner using a 16-channel knee coil at Brigham and Women’s Hospital, with MPRAGE sequences at 0.5 mm isotropic resolution. Additional acquisition details are provided in [32].

Retinotopic mapping was conducted using standard traveling-wave paradigms with flickering checker-board stimuli. Polar angle mapping utilized a rotating wedge stimulus, while eccentricity mapping employed an expanding or contracting annulus, both centered on a fixation point. Polar angle and eccentricity maps were projected onto each monkey’s reconstructed cortical surfaces. Borders between visual maps were delineated by reversals in polar angle representation at or near the horizontal, upper vertical, and/or lower vertical meridian. Full methodological details are available in prior publication [8]. For each monkey, retinotopic measurements were obtained separately for the left and right hemispheres, yielding a total of 12 hemispheres. Cortical surfaces were parcellated into nodes of approximately 1 mm^2^, and analyses were restricted to visual areas V1–V4. Each node was associated with its cortical spatial coordinates and a two-dimensional retinotopic tuning vector derived from fMRI responses. Individual hemisphere datasets are denoted using individual identifiers (M1–M6) and hemisphere labels (LH, RH), where LH and RH refer to the left and right hemispheres, respectively. Primary analyses were performed on the NMT population average template surface [47].

### Cortical Surface Geometry and Distance Metrics

The growth model was based on estimates of interaction strength between cortical locations, which depend on the distances separating them. Local intra-areal connectivity within gray matter tracks closely with distance along the cortical surface, while inter-areal projections traverse white matter pathways. Methods to accurately measure distances along these inter-areal fiber tracts are not established; however, relative distances along white matter pathways largely reflect relative distances along the cortical surface. We therefore used cortical surface distance as a proxy for the spatial constraints governing both local and long-range interactions in the growth model.

Cortical surface distances were computed pairwise between all surface nodes within a region of visual cortex encompassing areas V1, V2, V3, and V4. For each monkey and hemisphere, geodesic distances were computed along both the pial and smoothed white matter surface reconstructions using AFNI’s SurfDist. Distances computed on the pial and white matter surfaces were averaged to obtain a single cortical surface distance estimate for each node pair, providing a robust measure that accounts for folding-related differences between surface representations.

To obtain a uniformly sampled representation of cortex while preserving its intrinsic geometry, three-dimensional cortical coordinates were embedded into two dimensions using multidimensional scaling (MDS). The MDS embedding was computed using the node-wise cortical surface distance matrix and preserved relative distances between node pairs, with Spearman correlations between cortical surface distances and Euclidean distances in the 2D embedding exceeding 0.99. This distance-preserving embedding provides a flattened representation of the cortical sheet that retains intrinsic spatial relationships imposed by cortical geometry.

The embedding was resampled onto a uniformly spaced grid and the retinotopic tuning values were smoothed and interpolated using Matérn-kernel Gaussian process regression (*ν* = 2.5, length scale = 1.0, noise level = 0.1), with the two coordinate dimensions fit separately [48]. Each grid location was treated as a node in the network, forming the substrate for subsequent growth simulations. The resulting network comprised 3,486 nodes in NMT LH (1,334 in primary visual cortex and 2,152 in higher visual areas) and 3,238 nodes in NMT RH (1,110 in primary visual cortex and 2,128 in higher visual areas). Each node inherits both its position along the cortical surface and its retinotopic tuning from the underlying imaging data.

### Activity Correlation

Pairwise activity correlations between nodes were computed as a function of physical distance on the two-dimensional embedding of cortical locations. For each node *u*, distances to other nodes were calculated relative to a common foveal reference point. First, two orthogonal directions were defined at node *u*: a radial direction pointing from the fovea toward node *u*, and a tangential direction orthogonal to it. The distance between node *u* and another node *v* was then computed along the radial and tangential directions, denoted *d*_R_(*u, v*) and *d*_T_(*u, v*) respectively. To calculate the pairwise activity correlation between nodes *u* and *v*, the two distance components were combined into an exponential similarity kernel,

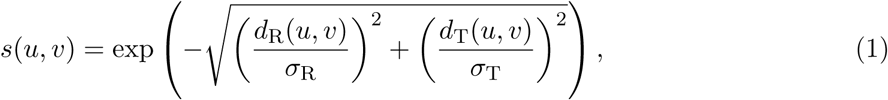

where *σ*_R_ and *σ*_T_ are parameters that determine the spatial extent of correlation along the radial and tangential directions, respectively. This parameterization was chosen to permit, but not require, directional bias in local interactions, because empirical studies in visual cortex indicate that local cortical spread can deviate from isotropy on the cortical sheet [49–51]. Heat maps of this similarity kernel for different values of *σ*_R_ and *σ*_T_ are shown in Fig. S1.

Using this fixed kernel, the net activity correlation between a V1 node *u* and an extrastriate node *v* was defined as

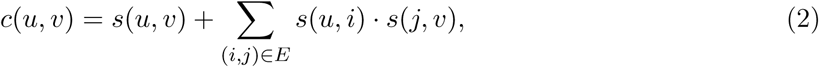

where *E* denotes the current set of directed edges. The second term in Eq. 2 captures indirect correlations across the network [16].

As a comparison, we also considered a simplified formulation of the similarity kernel in Eq. 1. Instead of separate radial and tangential components, we used the Euclidean distance between *u* and *v* to define the kernel. Specifically, we set *σ*_R_ = *σ*_T_ in Eq. 1, reducing the equation to *s*(*u, v*) = *exp*(−*d/σ*), where *d* is the Euclidean distance between *u* and *v*, and *σ* is the only parameter of the model (Supplementary Materials, Sec. G). Despite the reduced formulation, this single-parameter model produced qualitatively similar retinotopic organization (Fig. S7), indicating that the emergence of map structure does not depend critically on directional bias in local interactions.

### Degree-dependent Competition

This term captures competition among V1 nodes to form connections to extrastriate nodes. As the out-degree of a V1 node increases during network growth, its competitive weight decreases according to an inverse-degree function.

For a V1 node *u* with out-degree *k_u_*, the competition weight was defined as

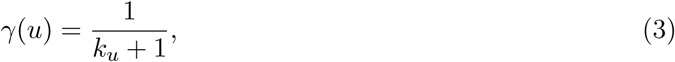

which is a monotonically decreasing and strictly positive function of degree. This formulation assigns lower competitive strength to highly connected nodes while ensuring that even the most connected nodes retain a nonzero capacity for further connections.

### Network Growth

The network grew by sequentially adding directed edges from nodes in V1 to nodes in extrastriate cortex, forming a directed bipartite architecture starting from an initially unconnected graph. At each growth iteration, an affinity score was computed between each V1 node and each unconnected extrastriate node (i.e. an extrastriate node that has not yet connected to the bipartite architecture),

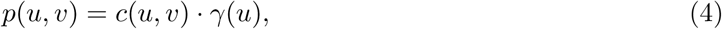

where *p*(*u, v*) denotes the affinity score between V1 node *u* and extrastriate node *v*, *c*(*u, v*) is the activity correlation between the two nodes (Eq. 2), and *γ*(*u*) is the competitive weight of node *u* (Eq. 3). The score quantified node *u*’s propensity to connect to node *v* [16]. The activity correlation term in Eq. 4 models a Hebbian mechanism, whereby correlated activity promotes the formation of new connections. After computing the affinities between each V1 node and each unconnected extrastriate node, all unconnected extrastriate nodes were ranked by their maximal affinities.

At each growth step, the single extrastriate node with the highest maximal affinity was selected as the target, and a directed edge was added from the V1 node for which it had the highest affinity. After each edge addition, the activity correlation and degree-dependent competition terms were updated. In the main experiments, nodes were incorporated sequentially in this manner. We also evaluated a batched growth variant in which extrastriate nodes were added in fixed-order groups to assess the robustness of the resulting organization to growth ordering (Fig. S4; Supplementary Materials, Sec. D).

### Parameter Settings

The model has only two free parameters: the radial and tangential components of the activity correlation kernel (*σ*_R_ and *σ*_T_ respectively in Eq. 1), which determine the spatial extent of activity correlations on the cortical surface. These parameters were tuned to match the retinotopic tuning values observed in the NMT dataset and were held fixed across the individual datasets. The tuned parameters were (*σ*_R_*, σ*_T_) = (1.30, 2.20) for both hemispheres.

We characterized the spatial spread of activity specified by the tuned parameters by examining how activity correlations decay as a function of geodesic cortical distance between V1 nodes (Fig. S3; see

Supplementary Materials, Sec. C). The model exhibits a consistent exponential decay of correlation with distance, with a characteristic length scale of approximately 1.86 mm across hemispheres. Notably, correlations plateau at distances of ∼5 mm, consistent with empirical measurements of distance-dependent response correlations in visual cortex [52].

### Robustness to Parameter Variations

In supplementary analyses, we evaluated the robustness of our results to the two parameters of our model (*σ*_R_ and *σ*_T_ in Eq. 1) by performing a grid search over these parameters (Supplementary Materials, Sec. B). Across a broad range of parameter values, the resulting maps remained qualitatively similar, preserving key features of retinotopic organization such as smooth phase progression and consistent areal structure (Fig. S2). Map structure degraded outside this parameter regime.

### Quantitative Comparison of Predicted and Empirical Maps

Model predictions were compared with empirically measured retinotopic maps using Spearman’s rank correlation, computed separately for polar angle and eccentricity. Analyses were restricted to extrastriate nodes (V2–V4) to assess retinotopic correspondence beyond the V1 anchor.

Polar angle phase was defined as the two-argument arctangent of the V1-anchored tuning vector, and eccentricity as the corresponding radial distance in this anchored coordinate system. Polar angle and eccentricity correlations were examined both individually and as a combined measure obtained by averaging the two values.

Retinotopic error was expressed in visual degrees by converting retinotopic coordinates from polar (*r, θ*) to Cartesian (*x, y*) space, with eccentricity normalized to 10^◦^. For each extrastriate node *v*, node-level error was defined as

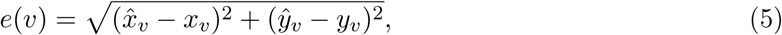

where (*x̂_v_*, *ŷ_v_*) and (*x_v_, y_v_*) denote predicted and empirical coordinates, respectively. Node-level errors were mapped onto the cortical surface to visualize spatial patterns of retinotopic correspondence. For each hemisphere, node-level errors were averaged, and the mean of LH and RH values was taken as the individual error.

### Angular Phase Analysis and Boundary Detection

To quantify the periodic structure of polar angle representations, retinotopic tuning coordinates were analyzed in the V1-anchored polar coordinate system described above. Nodes were then sorted by V1-based eccentricity and grouped into quantile-defined eccentricity bands. The most foveal 10% of nodes (corresponding to eccentricities within ∼ 1^◦^) were excluded because polar angle representations near the fovea are difficult to resolve with fMRI. The remaining nodes were binned into successive ∼ 10% eccentricity bands, with the final band slightly larger to include all remaining nodes.

For each eccentricity-band group, nodes were ordered by their angular position around the foveal center in a two-dimensional cortical embedding, and the largest contiguous V1 arc in this angular ordering was identified. The two endpoints of this V1 arc define two V1 border endpoints that correspond to the dorsal and ventral sides of the map. Using a precomputed geodesic distance matrix on the cortical surface, each extrastriate node (V2–V4) in the band group was assigned a signed cortical distance (in mm) to the nearer of the two V1 border endpoints, yielding separate distance axes for the dorsal and ventral sides. Cortical distance profiles were then computed using 1 mm-wide distance bins (bin size = 1.0 mm) by averaging phase values within each bin, separately for dorsal and ventral sides, and aggregating across eccentricity-band groups. Phase–distance agreement between empirical and predicted profiles was quantified by Spearman correlation computed over the binned mean profiles.

Boundary locations (V2/V3 and V3/V4) were operationally defined as reversal points (local extrema) in the phase-versus-distance profiles. For each hemisphere, extrema were identified separately on the dorsal and ventral profiles, yielding four boundary markers in total (two per side). Boundary positions were recorded as distances (mm) from the corresponding V1 border endpoint. To quantify relative area extent, node counts were computed in two ways: empirical area counts were obtained by summing the true anatomical area labels (V2/V3/V4) for all extrastriate nodes included in the eccentricity-band analysis, whereas predicted area counts were derived by assigning each extrastriate node to V2, V3, or V4 based on whether its absolute cortical distance from the V1 border fell below the predicted V2/V3 boundary distance, between the predicted V2/V3 and V3/V4 boundary distances, or beyond the predicted V3/V4 boundary distance (computed separately for dorsal and ventral sides within each hemisphere).

### Rotation Analysis

To assess whether correspondence between predicted and empirical retinotopy reflects specific alignment of retinotopic organization or generic mirror-reversing structure, V1 polar angle phase values were systematically rotated by fixed offsets (0^◦^, 45^◦^, 90^◦^, 135^◦^) prior to propagation through the model. Rotations were implemented as additive phase offsets applied to V1 polar angle values, with phase values wrapped to the original angular range after rotation.

For each rotation, extrastriate polar angle phase predictions were recomputed by applying the fixed V1-to-extrastriate projection weights obtained from the unrotated growth model; growth dynamics and connectivity were not rerun. Eccentricity values and all model parameters were held fixed. Spearman correlations and mean retinotopic error were computed between rotated predictions and empirically measured retinotopy, restricted to extrastriate nodes and using the same circular and normalization procedures as in the primary quantitative analyses.

### Cross-monkey Generalization Analysis

To evaluate the relative contribution of individual cortical geometry and V1 tuning statistics across monkeys, the growth model was applied to each monkey’s native cortical surface geometry while systematically substituting V1 retinotopic tuning from other monkeys.

For each target monkey, growth was performed on that monkey’s native cortical surface geometry using identical growth parameters. Model predictions were generated under two conditions: (1) using monkey-matched V1 retinotopic tuning and (2) using V1 tuning transferred from each of the remaining five monkeys, with transfer performed between homologous hemispheres.

Retinotopic correspondence was quantified using Spearman correlation (*ρ*) between model predictions and empirically measured polar angle maps across extrastriate nodes (V2–V4). For each monkey, retinotopic correspondence was summarized across hemispheres, and cross-monkey correspondence was summarized across the five transfer conditions. A two-sided sign test assessed whether within-monkey predictions systematically outperformed cross-monkey predictions. Under the null hypothesis that within- and cross-monkey predictions are equally likely to yield higher correspondence for each monkey, the two-sided probability that all six monkeys show the same direction of difference is 2 × (1*/*2^6^) = 1*/*32 ≈ 0.0312.

### Statistical Analyses

Statistical analyses quantified correspondence between model-predicted and empirically measured retinotopic organization and assessed monkey specificity of model predictions.

Agreement between predicted and empirical maps was quantified using Spearman correlation, computed separately for polar angle and eccentricity and restricted to extrastriate nodes (V2–V4).

Retinotopic error was quantified in visual degrees as the Euclidean distance between predicted and empirical retinotopic coordinates (Eq. 5). Node-level errors were averaged within each hemisphere, and the mean of left and right hemisphere values was taken as the subject-level error. Spatial patterns of retinotopic correspondence were assessed by mapping node-level errors onto the cortical surface.

Individual specificity in cross-monkey generalization analyses was evaluated using a two-sided sign-flip test comparing within-monkey and cross-monkey retinotopic correspondence across monkeys. No parametric assumptions were made about error distributions, and no correction for multiple comparisons was applied because each statistical test addressed a distinct, predefined analysis objective.

## Code Availability

Code implementing the growth model is available at https://github.com/ hhyy0401/vc_growth.

## Supplementary Materials

### A. Visualization of Activity Correlations

Activity correlations in the model depend on the similarity kernel in Eq. 1 of the main text. The kernel consists of two parameters *σ*_R_ and *σ*_T_. Fig. S1 shows example kernels obtained by varying these two parameters. When *σ*_R_ = *σ*_T_, the kernel is isotropic, corresponding to a symmetric fall-off of activity with distance. Increasing both parameters equally preserves this isotropy while broadening the overall spatial extent of correlations (top row of Fig. S1). Increasing the tangential parameter *σ*_T_ elongates the kernel along iso-eccentricity directions, corresponding to correlations across a broader range of polar angles. In contrast, increasing the radial parameter *σ*_R_ elongates the kernel along the radial direction, corresponding to correlations across eccentricities. These examples illustrate how the relative magnitudes of *σ*_R_ and *σ*_T_ determine the spatial spread of correlations.

**Figure S1:**
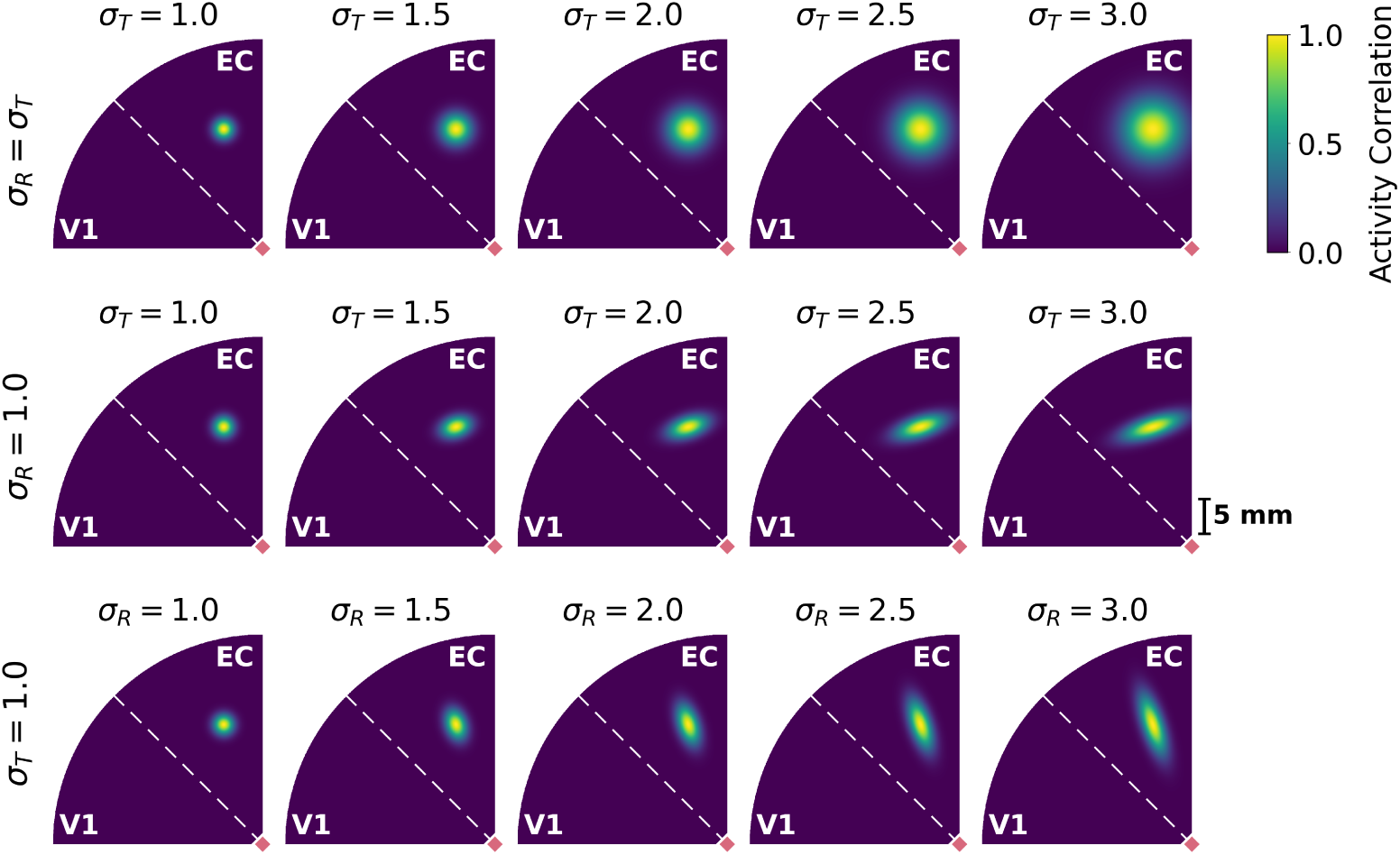
Activity correlation kernels for different values of the radial and tangential parameters. The kernel is centered at a reference location (yellow) in extrastriate cortex (EC). The fovea is indicated in red. The top row shows isotropic kernels obtained by increasing both parameters equally (*σ*_R_ = *σ*_T_), the middle row shows kernels obtained by varying the tangential parameter *σ*_T_ while fixing *σ*_R_ = 1.0, and the bottom row shows kernels obtained by varying the radial parameter *σ*_R_ while fixing *σ*_T_ = 1.0. The isotropic case produces a circular kernel, whereas increasing only one parameter produces anisotropic kernels elongated along either the tangential or radial direction.

### B. Robustness to Kernel Parameters

To evaluate the sensitivity of the model to the kernel parameters, we performed a grid search over *σ*_R_ and *σ*_T_. The radial parameter *σ*_R_ was varied from 0.5 to 2.5 in increments of 0.1 (21 values), and the tangential parameter *σ*_T_ from 0.5 to 2.5 in increments of 0.1 (21 values), yielding 441 parameter combinations per hemisphere. For each combination, we computed the mean retinotopic error on extrastriate nodes.

Figure S2A shows the resulting error landscape. To explore the region of reliable performance, we estimated a boundary separating near-optimal and degraded regimes. Specifically, for each value of *σ*_R_, we identified parameter combinations with the lowest 20% and highest 20% retinotopic error within that column. These labels were used to train a Gaussian Process Classifier with a radial basis function kernel [53]. The resulting classifier estimates the probability that a parameter combination belongs to the high-error regime. The boundary shown in Fig. S2A corresponds to the contour where the classifier assigns equal probability to the near-optimal and high-error regimes. At this contour, a parameter combination is equally likely to belong to the near-optimal or degraded regime, and thus the curve represents the transition between parameter regions that yield low versus high retinotopic error.

**Figure S2:**
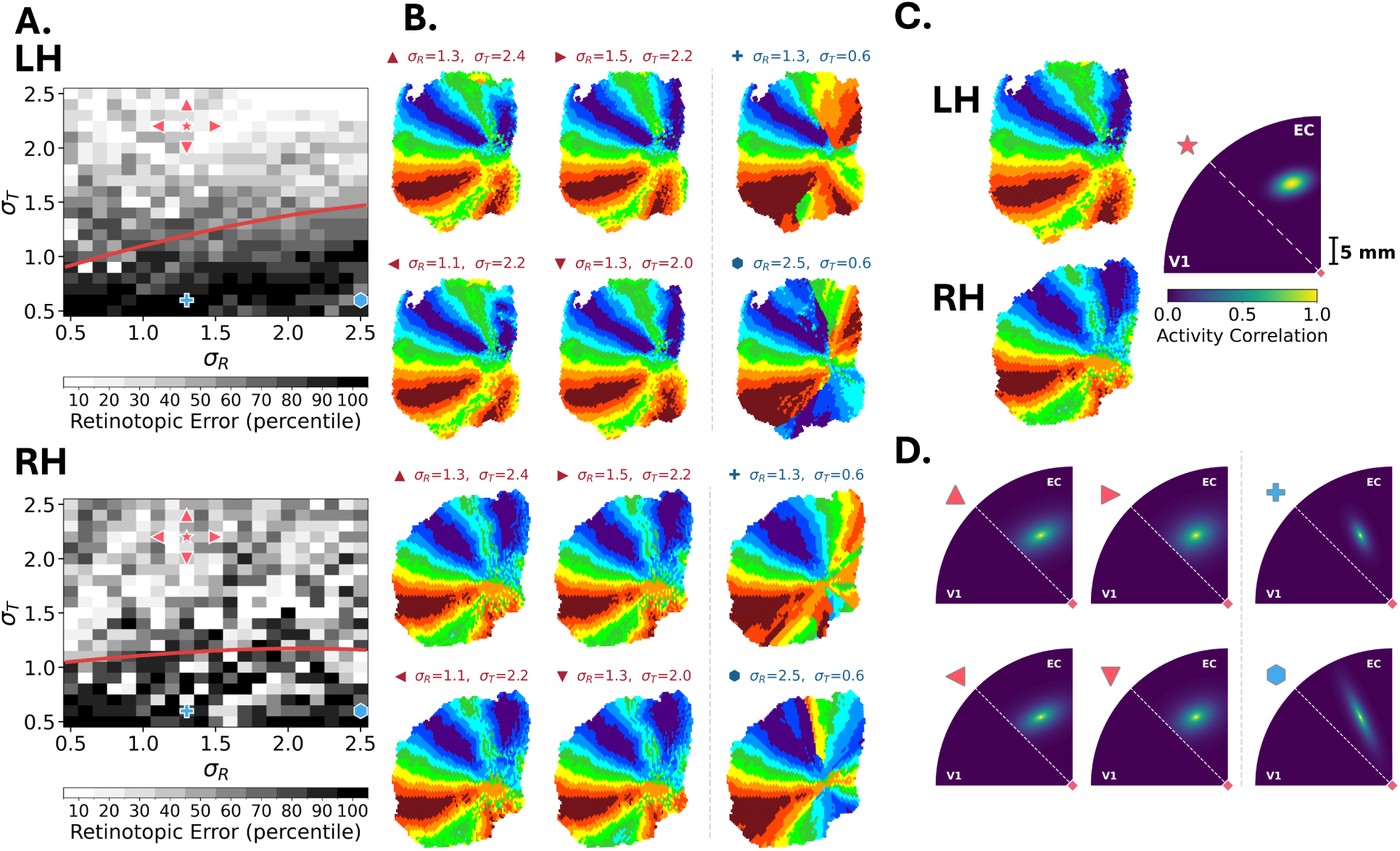
Robustness of retinotopic map formation to the kernel parameters (top: NMT LH; bottom: NMT RH). **A.** Retinotopic error across a grid search over *σ*_R_ and *σ*_T_ (0.5–2.5, step 0.1). Lighter shades indicate lower error, whereas darker shades indicate higher error. The red curve marks the boundary separating near-optimal and degraded parameter regimes. Parameter combinations on the near-optimal side of this boundary yield low retinotopic error, whereas those beyond it produce substantially worse predictions. Marker symbols correspond to the parameter settings visualized in panels B–D; red markers denote near-optimal settings and blue markers denote outlier settings outside the boundary. **B.** Six predicted polar-angle maps for representative parameter settings are shown. The first four panels correspond to parameter settings near the optimal region (*▴*, *▶*, *◀*, *▾*), for which retinotopic organization remains stable. The rightmost column corresponds to two outlier parameter settings outside the boundary (+, ⬢), where retinotopic organization deteriorates. **C.** Predicted polar-angle maps for LH and RH using the optimal parameter setting (*σ_R_* = 1.3*, σ_T_* = 2.2, denoted as *⋆* in panel A), together with the corresponding elliptical correlation kernel (visualized as in Fig. S1). **D.** Six predicted elliptical correlation kernels corresponding to the marked parameter settings in panel B.

**Figure S3:**
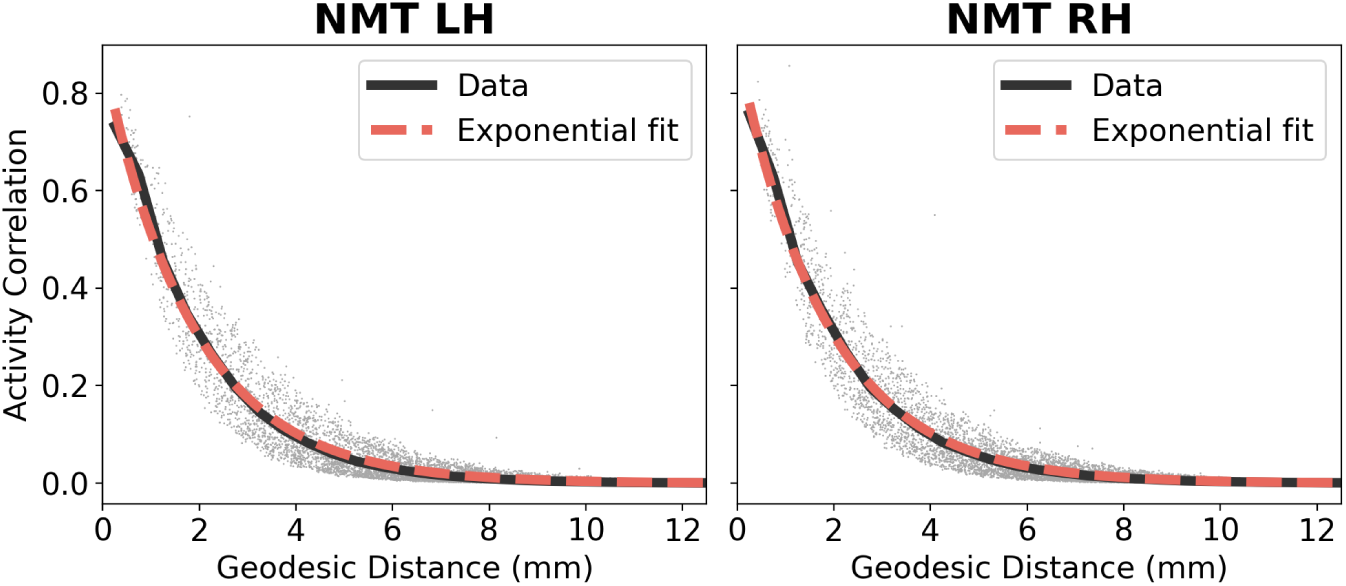
Spatial decay of activity correlations implied by the similarity kernel. Kernel similarity between pairs of V1 nodes is plotted as a function of geodesic cortical distance along the cortical surface for NMT LH and RH, respectively. Gray dots indicate individual node pairs (subsampled for visualization), solid dark curves show binned means, and dashed red curves show exponential fits of the form *K*(*d*) = *A* exp(−*d/σ*).

Predicted polar-angle maps for representative parameter settings are shown in Fig. S2B–C (C: optimal parameters). Maps remain qualitatively stable for parameter values surrounding the optimal region, but begin to deteriorate outside the boundary, where smooth gradients and mirror-reversing organization are disrupted.

Fig. S2C–D (C: optimal parameters) visualizes correlation kernels across representative values of *σ*_R_ and *σ*_T_ used in Fig. S2A-B. Retinotopic error showed a modest tendency to be lower when the tangential parameter exceeded the radial parameter (*σ*_T_ *> σ*_R_), producing kernels that extend along the tangential axis and support interactions across a broader range of polar angles while preserving eccentricity structure. This tendency toward tangential elongation is broadly consistent with fMRI studies reporting anisotropic local correlation structure in early visual cortex with respect to retinotopic coordinates [54–56]. In contrast, when *σ*_T_ *< σ*_R_ the kernel becomes radially biased, reducing angular coverage and degrading retinotopic organization. When *σ*_T_ = *σ*_R_, the kernel reduces to an isotropic (Euclidean) distance kernel.

### C. Spatial Decay of Activity Correlations

Here we examine how predicted activity correlations between V1 nodes vary as a function of geodesic cortical distance along the cortical surface (Fig. S3). Geodesic distances between all pairs of V1 nodes were measured on the cortical surface mesh.

For each pair of nodes (*i, j*), the similarity kernel *s*(*i, j*) was computed with parameters fixed to the optimal values (*σ*_R_ = 1.30, *σ*_T_ = 2.20) used in the main text. This kernel is our model’s prediction of the spatial spread of activity correlations in the macaque brain.

The kernel values were grouped into 0.5 mm distance bins to summarize how activity correlation varies with cortical distance. We then fit an exponential function *K*(*d*) = *A* exp(−*d/σ*) to the binned means. The fitted parameters were *A* = 0.88, *σ* = 1.86 mm for NMT LH and *A* = 0.89, *σ* = 1.86 mm for NMT RH, indicating a characteristic decay length of *σ* ≈ 1.86 mm (the distance at which correlation falls to *A/e*). Notably, activity correlations bottom out at ∼5 mm in both hemispheres, which is broadly consistent with empirical observations of response correlation decay as a function of cortical distance reported in prior work [52].

**Figure S4:**
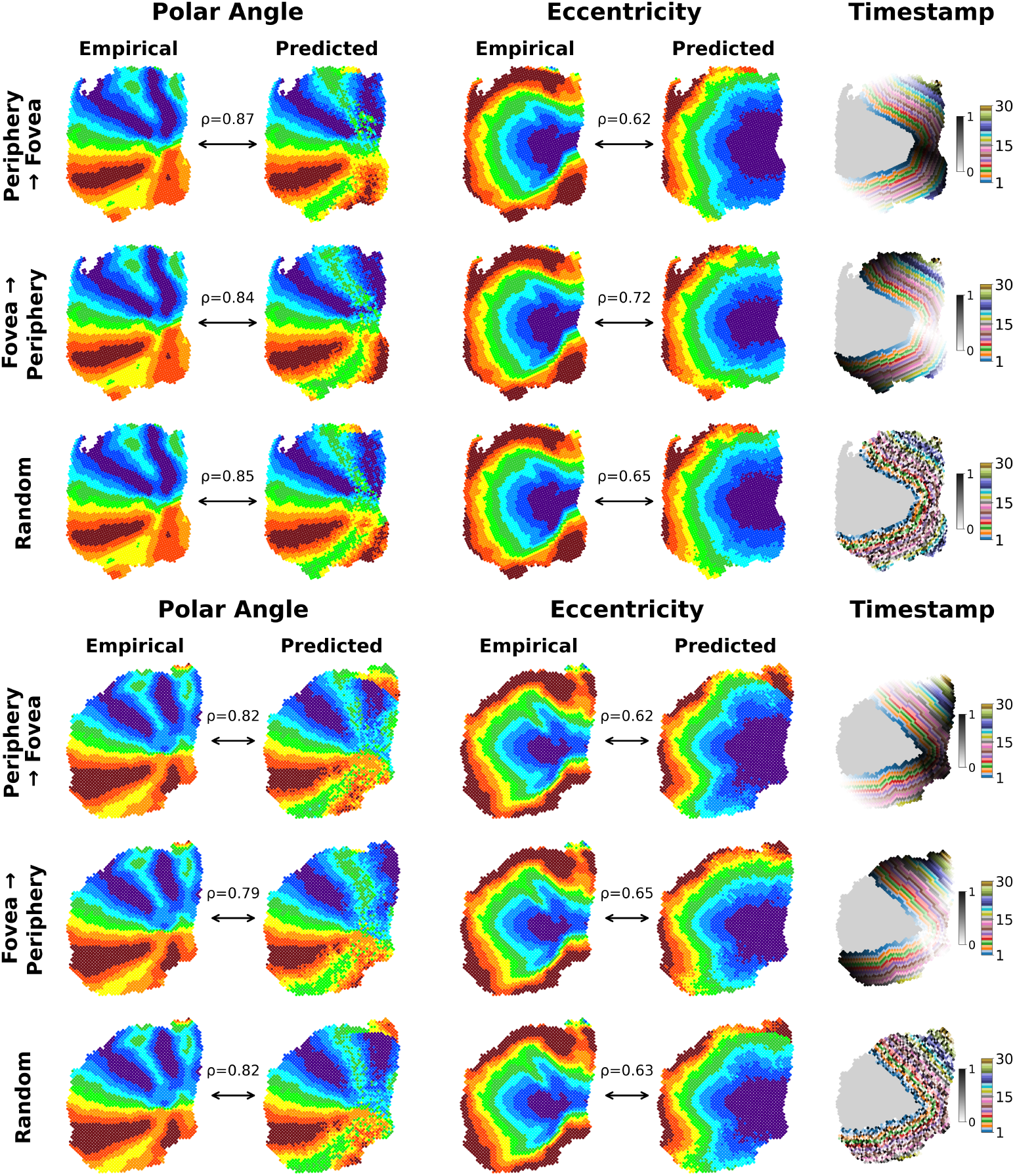
Effects of batch-ordering schemes on retinotopic map formation in LH (top) and RH (bottom). Extrastriate nodes were grouped into 30 distance bins and incorporated in three orders: periphery→fovea, fovea→periphery, and random (rows). Polar angle maps show strong agreement across conditions (LH: *ρ* = 0.84–0.87; RH: *ρ* = 0.79–0.82), preserving mirror-reversing structure. Eccentricity maps remain smooth and radial with stable correlations (LH: *ρ* = 0.62–0.72; RH: *ρ* = 0.62–0.65). Timestamp maps (rightmost column) indicate the order of node incorporation, where colors denote distance bins (1–30), and within each bin values range from 0 (lighter, white-mixed tones) to 1 (darker, black-mixed tones). Despite different growth trajectories, final map organization is consistent across orderings.

### D. Robustness to Growth Order

To assess whether the emergence of retinotopic organization depends on the temporal sequence of cortical growth, we systematically varied the order in which extrastriate nodes were incorporated into the model. Nodes were grouped into 30 bins based on cortical distance from the foveal anchor, computed using Euclidean distance. The same parameter values (*σ*_R_ and *σ*_T_) used in the main experiment were retained. Three batch-ordering schemes were tested: progression from periphery to fovea, from fovea to periphery, and a random ordering. Within each batch, node selection followed the same affinity-based rule as in the primary model, ensuring that only the ordering was manipulated.

Across all ordering conditions, the model produced highly consistent retinotopic maps (Fig. S4). Both polar angle and eccentricity organization remained stable, exhibiting strong agreement between empirical and predicted maps. In NMT LH (top), polar angle correlations were identical across all conditions (*ρ* = 0.84–0.87), while eccentricity correlations ranged from *ρ* = 0.62 to 0.72. In NMT RH (bottom), correlations were slightly lower but remained robust, with polar angle values of *ρ* = 0.79–0.82, and eccentricity values ranging from *ρ* = 0.62 to 0.65. Despite minor variability, no systematic shifts were observed in map orientation, boundary placement, or topographic structure. These findings indicate that the emergence of retinotopic structure is invariant to growth order and instead arises from local distance-dependent interactions and competitive dynamics.

### E. Hierarchical Growth Model

In the primary model, cortical organization emerges through a directed bipartite growth process in which projections originate exclusively from V1 and extend to higher visual areas. This formulation reflects a minimal anchoring assumption, in which V1 provides the sole developmental reference frame for extrastriate organization. However, the visual cortex has hierarchical interareal connectivity, raising the question of whether a hierarchical developmental process would alter map organization.

To evaluate this possibility, we implemented a hierarchical model in which development unfolds in sequential stages across putative visual areas. Growth is initiated from V1 as in the model presented in the main text. Once most of V1 nodes establish outgoing connections, the subset of extrastriate nodes receiving these connections is treated as the source population for the next stage of growth. These nodes then form directed projections to a downstream population corresponding to the next visual area. This process is repeated iteratively, with transitions between stages occurring once 90% of nodes in the current source population establish outgoing connections. Apart from this staging rule, all components of the model, including the similarity kernel (Eq. 1) and affinity function (Eq. 4), were the same as the model in the main text, with *σ*_R_ and *σ*_T_ set to 1.40 and 2.20 in both hemispheres. We explored parameter settings specific to the hierarchical formulation to ensure a fair evaluation of its performance. Mechanistically, this hierarchical growth sequence can be instantiated through an attenuation factor [17] that suppresses cascading activity in the network, allowing earlier areas to establish outgoing connections before later areas form theirs.

Under this hierarchical formulation, the model produced retinotopic organization (Fig. S5) similar to that presented in the main text. Spearman’s *ρ* between model and data was 0.87 (LH) and 0.85 (RH) for polar angle, and 0.64 (LH) and 0.63 (RH) for eccentricity. These results indicate that hierarchical growth is not required to generate higher-order retinotopic organization, but is nonetheless compatible with the developmental mechanism presented in the main text. The results also imply that emergence of map structure is primarily determined by local interaction rules and cortical geometry, rather than by the developmental sequence of interareal connectivity.

**Figure S5:**
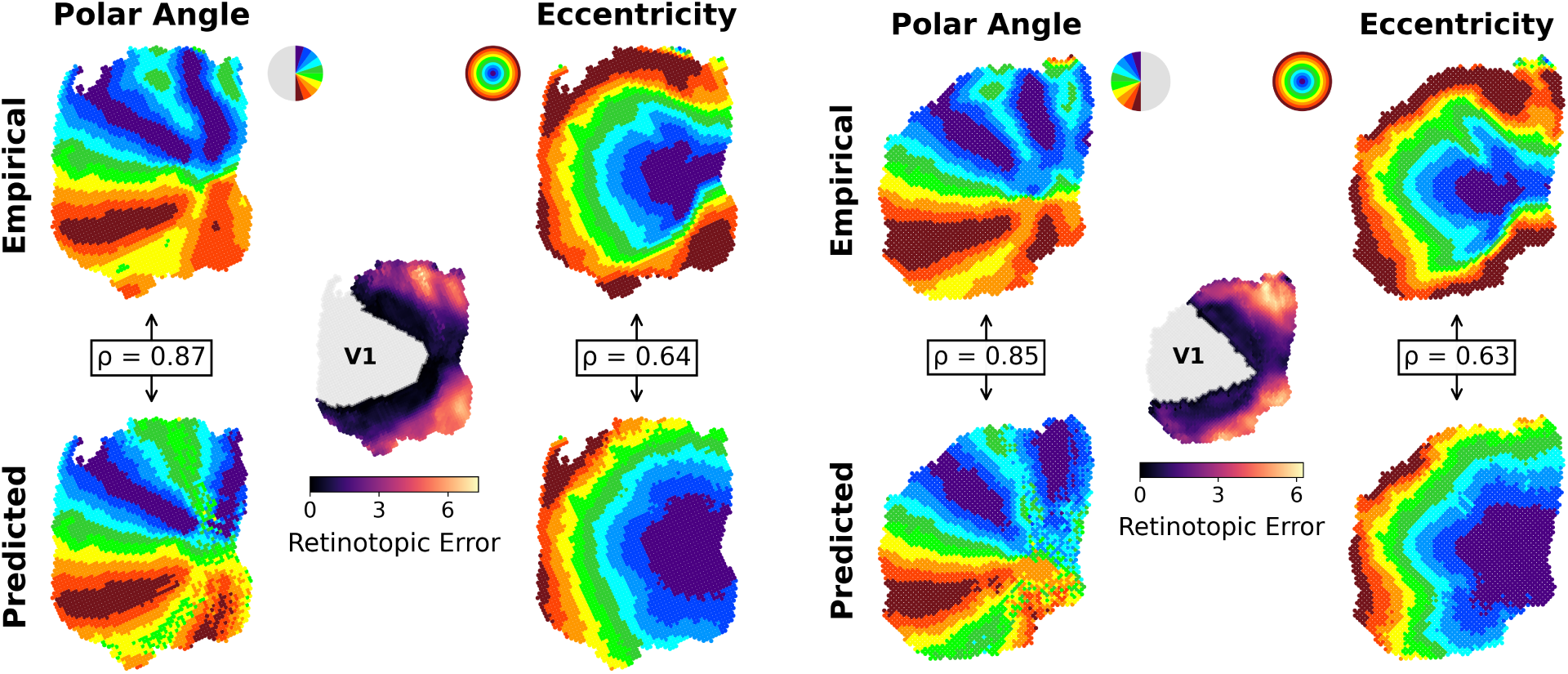
Hierarchical growth model predictions on the NMT cortical geometry (left: NMT LH, right: NMT RH). For NMT LH, non-V1 Spearman’s rank correlation is *ρ* = 0.87 for polar angle and *ρ* = 0.64 for eccentricity. Similar results are observed in NMT RH, with *ρ* = 0.85 for polar angle, *ρ* = 0.63 for eccentricity. The error maps show increasing error with distance from V1.

### F. Model predictions across individual cortical geometries in RH

Fig. S6 illustrates how the model generalizes across individual cortical geometries in the right hemisphere. Despite anatomical differences (Fig. S6A), the model captures consistent retinotopic organization with accurate phase progression and mirror reversals (Fig. S6B), and reproduces monkey-specific eccentricity gradients aligned with empirical maps (Fig. S6C). Together with the results of the main text, these plots indicate that the model robustly adapts to individual cortical geometries while preserving key topographic principles.

**Figure S6:**
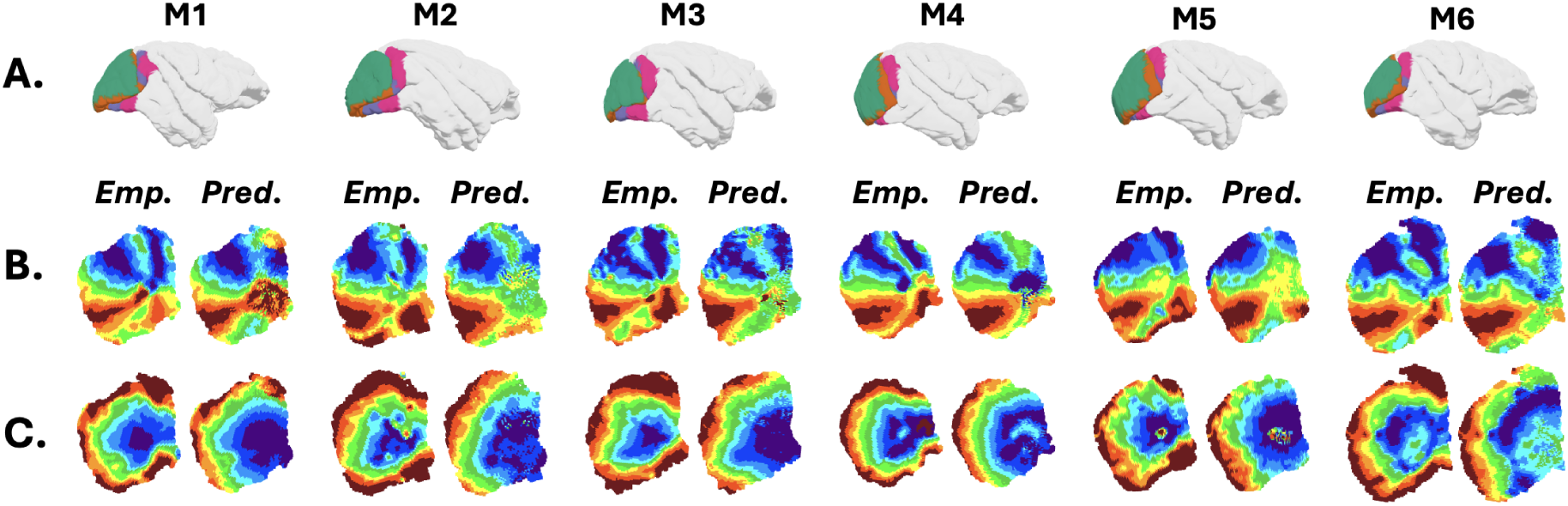
A. Three-dimensional cortical surface reconstructions for the right hemisphere in each dataset, with visual areas V1–V4 indicated by distinct colors, illustrating dataset-specific cortical geometry and areal layout used as the growth substrate. **B.** Retinotopic phase maps, comparing empirically measured maps (empirical) with model predictions (pred) for each monkey. Across datasets, higher visual areas show smooth phase progression within maps and consistent mirror reversals at map boundaries. **C.** Corresponding eccentricity maps showing the radial component of retinotopic organization. The model reproduces monkey-specific eccentricity gradients aligned with empirical measurements.

### G. Reduction to Euclidean Distance Formulation

In the primary model, activity correlations are defined using a decomposition of cortical distance into radial and tangential components (Eq. 1 of main text), introducing two parameters that separately control the decay of correlations along these dimensions. To assess map formation when the decays are equal in both dimensions, we replaced the radial–tangential decomposition with a single Euclidean distance formulation. Specifically, activity correlations were computed as a function of Euclidean distance *d*(*u, v*) in the two-dimensional cortical embedding, using an isotropic similarity kernel,

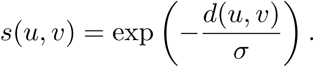

This model is equivalent to the primary formulation described in the main text, with *σ*_R_ = *σ*_T_. The parameter *σ* was tuned to match the macaque retinotopic data and was set to *σ* = 2.8.

This single-parameter model produced retinotopic maps similar to the two-parameter model (Fig. S7). Polar angle maps showed strong correspondence to empirical observations of mirror-reversing structure (NMT LH: *ρ* = 0.87, NMT RH: *ρ* = 0.85), while eccentricity maps exhibited moderate agreement (NMT LH: *ρ* = 0.61, NMT RH: *ρ* = 0.66). Overall spatial organization remained smooth, with errors increasing gradually with distance from V1.

These results indicate that retinotopic organization does not depend on anisotropic correlations, but instead emerges from a monotonic distance-dependent interaction rule.

**Figure S7:**
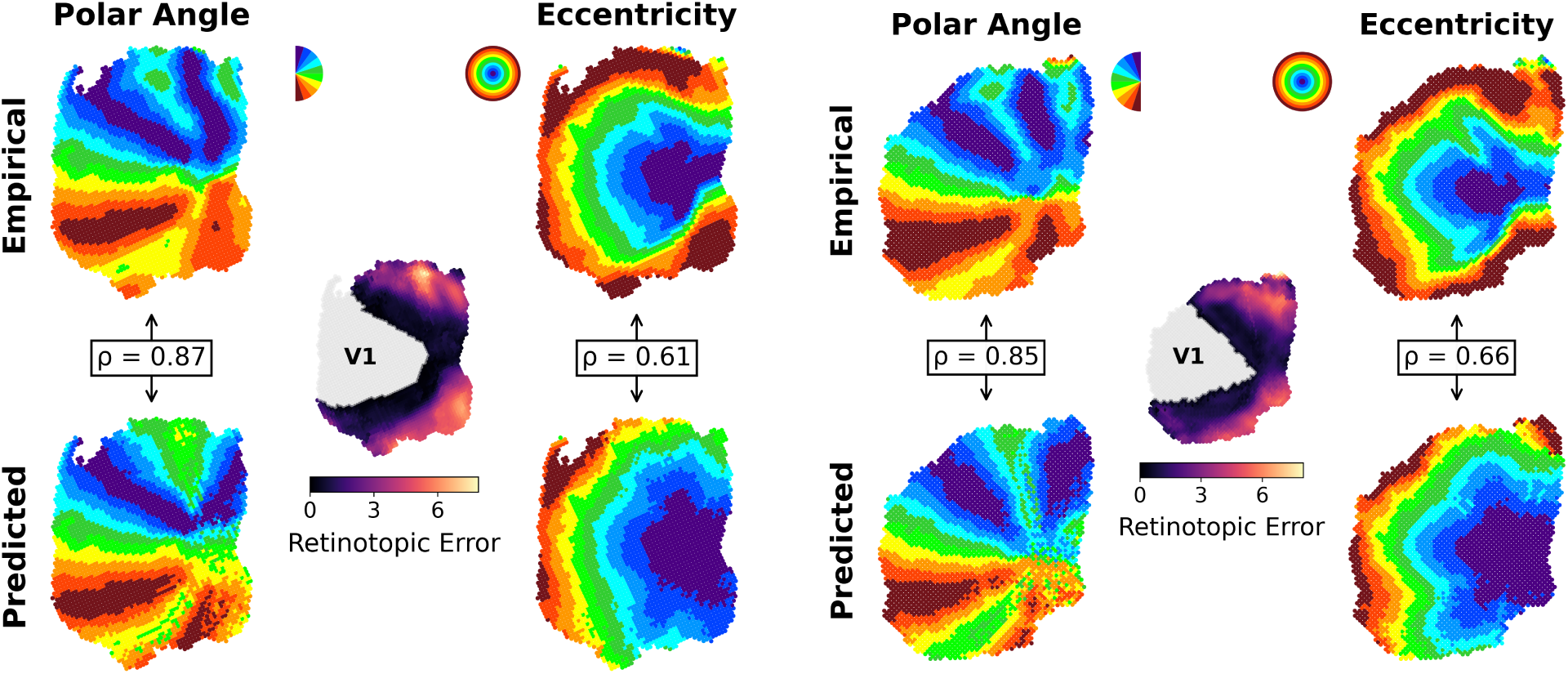
Retinotopic maps obtained from a single-parameter model using a Euclidean distance formulation (left: NMT LH, right: NMT RH). The same parameter setting was used for each hemisphere (*σ* = 2.8). Polar angle maps show strong agreement with empirical data (LH: *ρ* = 0.87, RH: *ρ* = 0.85), while eccentricity maps exhibit moderate correspondence (LH: *ρ* = 0.61, RH: *ρ* = 0.66). Overall topographic structure is preserved despite the simplified formulation.

